# Structural Aspects of Nipah Virus Glycoproteins using Predictive Protein Modeling

**DOI:** 10.64898/2026.09.02.748953

**Authors:** Urmi Roy

## Abstract

Nipah is a virus borne zoonotic disease that mostly occurs in Southeast Asian countries. Nipah virus (NiV) infection is potentially lethal as it can cause fatal encephalitis, and as of now, there are no vaccines or therapeutic treatments for this disease. The highly pathogenic NiV belongs to the Paramyxovirus family and the natural host of NiV is fruit bats of the *Pteropodidae* family. Though NiV has not been detected in the US, rather recently the Camp Hill virus (CHV) in shrew species was identified in Camp Hill, Alabama. This virus is characteristically similar to Nipah, and belongs to the *henipavirus* genus that include the Nipah and Hendra viruses.

NiV has two surface glycoproteins namely the attachment glycoprotein (NiVG) and the fusion glycoprotein (NiVF). Being associated with host cell attachment and fusion of viral membrane, these glycoproteins tend to facilitate the viral entry to the host cell.

The present paper describes the structures of the aforesaid viral glycoproteins based on the available database of protein structures, and examines if they have an evolutionary link to other viral glycoproteins. Certain similar glycoproteins found in different countries are also explored. The structure-function relationship for these two NiV glycoproteins are determined using time-based molecular dynamics (MD) simulation. The interfacial ligand-receptor interactions between NiV proteins with human receptor and monoclonal antibody, Fab 66, are examined. A primary motivation for these structural studies is that the results might potentially contribute to future therapeutic developments and drug designs to effectively combat NiV.

## Introduction

Outbreaks of Nipah disease are generally reported in South/Southeast Asia. This highly pathogenic lethal virus was first identified in Malaysia in 1998, and later reported in Bangladesh, India, Philippines, Singapore and other Southeast Asian countries [1]. According to WHO, several laboratory-confirmed cases of Nipah virus (NiV) infection were found in India in 2026. Though there are no other confirmed cases, but this proves that the pathogen has not been eradicated yet [2]. Henipavirus nipahense (Nipah virus; NiV) is the primary pathogen for causing this disease, is commonly transmitted by bats or pigs through direct contact via body-fluids or respiratory secretions. The disease is typically manifested in the form of respiratory disorder and encephalitis, which can spread from person to person. The natural host of NiV is fruit bats of the *Pteropodidae* family that includes Pteropus genus.

The highly pathogenic NiV belongs to the Paramyxoviruses group that also includes the Hendra virus (HeV). The *henipavirus* genus, within the family *Paramyxoviridae* contains some of the most pathogenic *Paramyxoviruses*, including NiV, HeV and Cedar virus (CedPV). Other examples of *paramyxoviruses* include parainfluenza and mumps viruses. Although the symptoms of Nipah often are mild, it can also lead to fatal encephalitis [3]. According to the Center for Disease Control (CDC), 40%-70% of Nipah infections can potentially turn to be lethal. The World Health Organization (WHO) has identified Nipah as a possible future epidemic while the CDC listed NiV as “Category C agent”. Both NiV and HeV are identified as Biosafety level (BSL) 4 organisms.

Though NiV has not been previously detected in the US, rather recently the Camp Hill virus (CHV) has been detected in the form of short-tailed shrew species in Camp Hill, Alabama [4]. This has been the first report of a *henipavirus* in North America. Although CHV has not been found so far in humans, it exhibits many similarities with NiV, and belongs to the *henipavirus* genus that include the Nipah and Hendra viruses. Although past occurrences of NiV have mostly been localized in Southeast Asia, Australia and some of the African countries, it is evident after the last corona pandemic that, any virus capable of spreading rapid infections can pose worldwide public health threat. Several years back, GHP-88309 a non-nucleoside inhibitor was identified for some of the Paramyxoviruses [5], but to date no effective treatments are available for this deadly Nipah pathogen.

The enveloped Nipah virus has a negative single-stranded RNA genome. Nipah virus contains six structural proteins namely nucleocapsid (N), phosphoprotein (P), matrix protein (M), fusion glycoprotein (F), receptor binding or attachment glycoprotein (G) and the RNA polymerase (L). On the surface of the Nipah virus two spike proteins are present, namely the attachment glycoprotein (NiVG) and the fusion glycoprotein (NiVF), which are associated, respectively, with the receptor (host) cell attachment and membrane fusion [6, 7]. Usually, the host cell attachment is mediated by the NiVG protein with the surface receptor ephrinB2 or B3. NiVF enables fusion between NiVF and the host membrane and thus, binds to the host membrane protein, which consequently facilitates viral entry. These two proteins NiVG and NiVF, would also serve as targets for drug/antibody therapy. In addition to the six major structural proteins, NiV contains three non-structural proteins V, W and C, all encoded by the phosphoprotein (P) gene. These non-structural proteins inhibit their host’s interferon (IFN) induction, and help to escape the host’s immune system to develop infection. It is useful to note in this context that unlike the cases of NiV and HeV, the conserved V protein is absent in CedV [8].

In a number of previous papers, the present author has elucidated the structure of several immune system proteins [9–11]. Some contemporary virus structures and the host-virus interactions have also been discussed [12, 13]. In this report, we will analyze the structure of the NiV and the time-based stability of the host receptor bound NiVG as well as the antibody bound NiVF.

This work utilizes the currently available protein structures [6, 7], and aims at performing a relatively detailed protein-protein structural analyses. A brief evolutionary analysis of NiVG and NiVF with similar viral proteins is also included. The investigation centers on computational structural virology including protein-structure analyses, MD simulations, and phylogenetics. MD simulations of such systems are subject to certain limitations such as force field inaccuracies, time-scale limitations, and lack of quantum mechanical information about electronic transitions and bond-transformations. At the same time the MD simulation approach for protein structural studies offer several advantages such as efficient modeling of the protein’s inherent dynamic behavior, gaining mechanistic insight about microscopic structural changes, accessing thermodynamic information (binding affinities and folding pathways), and utilizing classical trajectories that are unbiased from prior experimental biases. With these later attributes, predictive modeling based on artificial intelligence and machine learning can largely facilitate the tasks of predicting immunogenicity, epitope mapping, and developing strategies for antibody-based therapeutics. Additionally, these simulation-based virtual screenings are highly effective in saving cost and time for systematic investigations.

Since NiV is highly pathogenic and since there are no vaccines or therapeutics are available for NiV at this time, detailed structural evaluation of NiV protein can be quite useful to develop structure-based drug designs and targeted therapies. The evolutionary and phylogenetic analyses of the NiV protein is important to trace its mutants and future lineages. Such analyses are also useful to understand the viral epidemiology and to identify as well as build neutralizing antibodies and vaccine candidates.

## 2. Methods

### 2.1. Protein Structure

For the receptor bound Nipah virus structure, we have used the protein data bank (PDB) structure 2VSM.PDB [6]. 2VSM is the human cell surface receptor ephrinB2 bound to NiVG. For the antibody bound virus we have employed the 6T3F.PDB [7]. 6T3F is the NiVF glycoprotein bound monoclonal antibody (mAb) Fab66. While previous authors have described simulation studies of ephrinB2 bound NiVG, the present study is based on a different simulation scheme [14].

### 2.2. The Modeling Software and the Modeling Process

Nanoscale Molecular Dynamics (NAMD), Visual Molecular Dynamics (VMD) and QwikMD software packages are used for modeling [15–17]. Proteins time-based structural changes are identified using MD simulation [18]; to do this, the protein structure files (psf) were first generated using VMD, and the system energy was minimized in 2000 steps. After the minimization steps, the annealing process was applied for 0.24 ns and the temperature of the system was ramped up from 60 K to 300 K. The equilibration steps were for 0.04 ns and then the final MD simulation was run for 50 ns. All simulations were carried out in an implicit solvent system using the Generalized Born/solvent-accessible model. The temperature was kept at 300 K using Langevin dynamics. During the final run no atoms were constrained while the backbone was restrained during the annealing and equilibration processes. For further elucidation, simulations of these two PDB structures were performed using an explicit solvent system. The details of these simulations have been included in the Supplementary Information, SI.

### 2.3. The Protein Structure Modeling

2VSM is the NiVG (chain A) bound to the human cell surface receptor ephrinB2 (EFNB2; chain B) [6]. For MD simulation, we have used 2VSM with chains A and B. 6T3F.PDB is the X ray crystal structure of NiVF protein (F chain) with antigen-binding fragment (Fab66) light (L) and heavy (H) chains [7]. For MD simulation, we have employed the truncated form of the 6T3F.PDB, and used residues 50-285 for the NiVF (F chain); residues 1-111 and 10-112 were used for the L and H chains of FAB66 receptor, respectively. Altogether, two simulations were conducted for this study using implicit solvent models. For the first simulation the 2VSM structure, NiVG (chain A) bound EFNB2 (chain B) was taken, while the second run was conducted using the truncated form of the 6T3F.PDB. The ligand F (chain F) bound receptor Fab66 (L and H chains) was considered in the latter case. The resulting protein structures were generated by Discovery Studio [19].

The MD simulation approach is used here to analyze the physical movements of atoms and molecules using Newton’s laws of motion. Based on the advantages of this technique (noted in the Introduction Section) this approach is used here to monitor the real-time atomistic movements of the NiV protein structure, and to assess the stability of the system, while determining the protein’s binding affinities, structure-function relationships, protein-protein as well as protein-small molecule/drug interactions as functions of time. A primary goal of this work is to generate generating potentially useful contributions to the database that supports the field of computational drug design. The classification techniques of this simulation study include analyses of the structural metrics, namely Root Mean Square Deviation (RMSD), Root Mean Square Fluctuation (RMSF) and secondary conformational changes of the protein.

### 2.4 Phylogenetic analysis

The evolutionary relationship of NiVG, NiVF and other viral glycoproteins and fusion proteins have been analyzed herein by Molecular Evolutionary Genetics Analysis 7 (MEGA 7) [20]. The Bootstrap method was used to perform a test of phylogeny. This evolutionary history was inferred using the Neighbor-Joining method [21]. Any gaps between data points or missing data were deleted, and the number of bootstrap replication was kept at 200 [22]. The Poisson model was used to calculate the evolutionary distances [23]. Evolutionary analyses of the NiVG and NiVF from different subcontinents are presented in this paper (SI Figure S1-S2), and the associated phylogenetic trees are generated using MEGA.

### 2.5. Predictive Protein-Protein Network

The protein-protein interaction network was predicted using open-source software platform Cytoscape [24]. The UniProt ids for NiVG and NiVF used in these networks were Q9IH62 and Q9IH63 [25].

## 3. Results and Discussion

2VSM is an X-ray crystal structure at 1.80 Å resolution, while EFNB2 and EFNB3 are the two host cell receptors for NiVG. The chain A of 2VSM consists of NiVG viral glycoprotein. The chain B is the EFNB2 receptor protein of the host. 6T3F.PDB is another X-ray crystal structure with a 3.20 Å resolution. The chain A represents the F glycoprotein of NiV while the chains L and H represent the light and heavy chains of Fab66, respectively. Figure 1 displays the 3D structures of NiV bound receptor complex. Figure 1A represents the structure of 2VSM.PDB, where the NiVG viral glycoprotein (red) and host receptor protein EFNB2 (blue) are displayed. The inset shows the G-H loop (green) where the β chains G and H are shown in cyan and violet, respectively. Figure 1B depicts the complete 6T3F structure and Figure 1C shows the truncated form of 6T3F.PDB.

**Figure 1.**
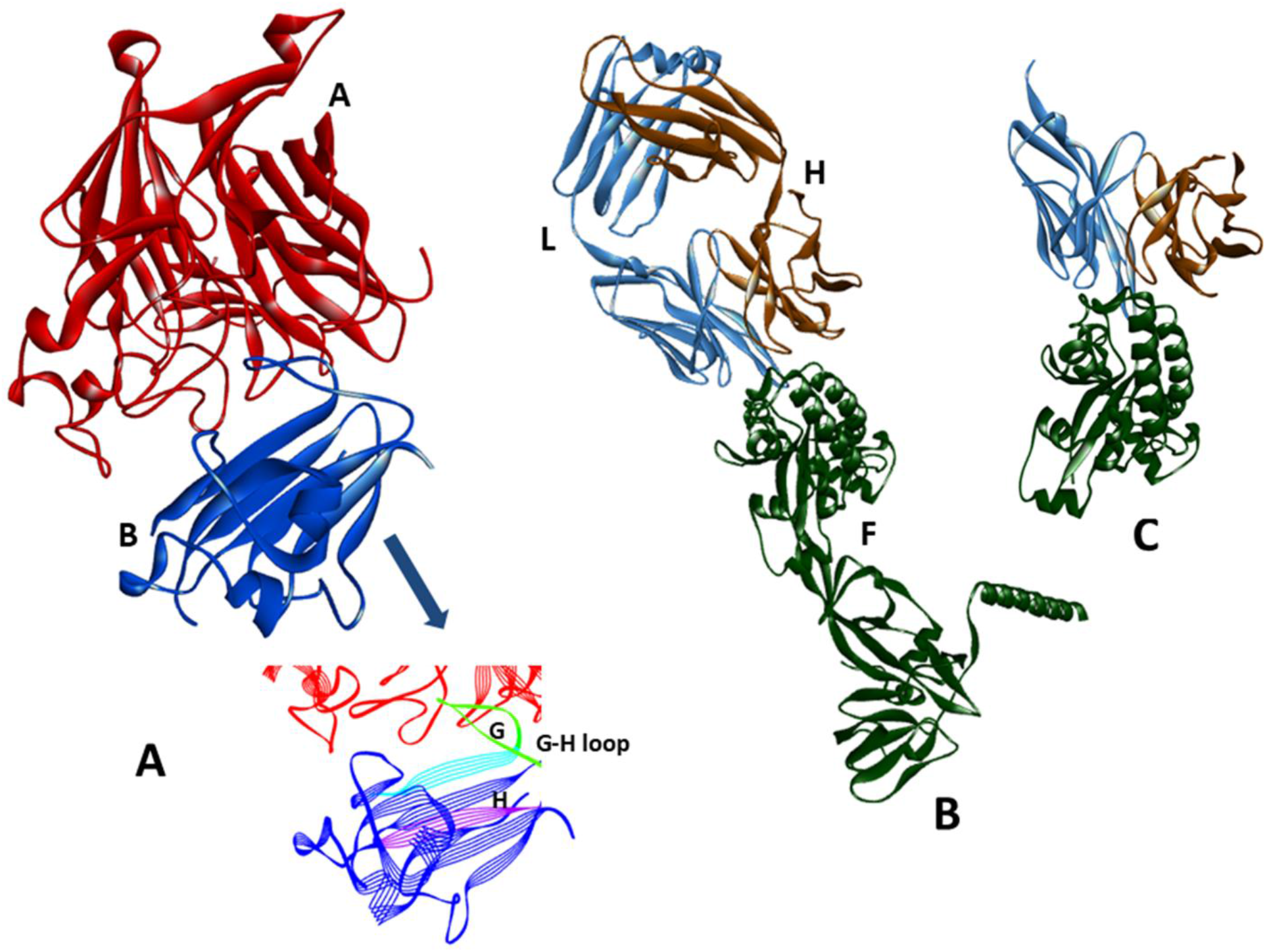
The 3D structure of A. 2VSM.PDB. Chain A of 2VSM represents NiVG viral glycoprotein (red) and chain B represents the host receptor protein EFNB2. Inset shows 2VSM: B where the G-H loop is colored in green. β chains G and H are colored in cyan and violet. B. The 3D structure of 6T3F.PDB. The NiV fusion protein is represent by F chain (green) and the FAB heavy and light chains are represented by L (Blue) and H (Brown) chains of 6T3F respectively. C. The truncated NiVF protein bound FAB66 receptor chains are represented here.

Figure 2A represents the evolutionary relationship of glycoproteins with 10 emerging zoonotic viruses, which corresponds to the most appropriate tree where the sum of branch lengths is 5.24. Here we have used UZQ22167, the most recent reported NiVG sequence from Figure S1 [26]. The viral glycoproteins/attachment G proteins/cell attachment protein sequences included in this chart are based on the following series of proteins: NiV G along with CedPV attachment G, Ghana virus G, Hendra virus G p, J virus attachment G, Melian virus cell attachment, Mojiang virus attachment G, Mount Mabu Lophuromys virus 1 G and 2 G and Paramyxoviridae sp. attachment G (partial) [8, 26–34]. The phylogenetic tree of 17 NiVG protein sequences reported from different countries are displayed in Figure S1.

**Figure 2.**
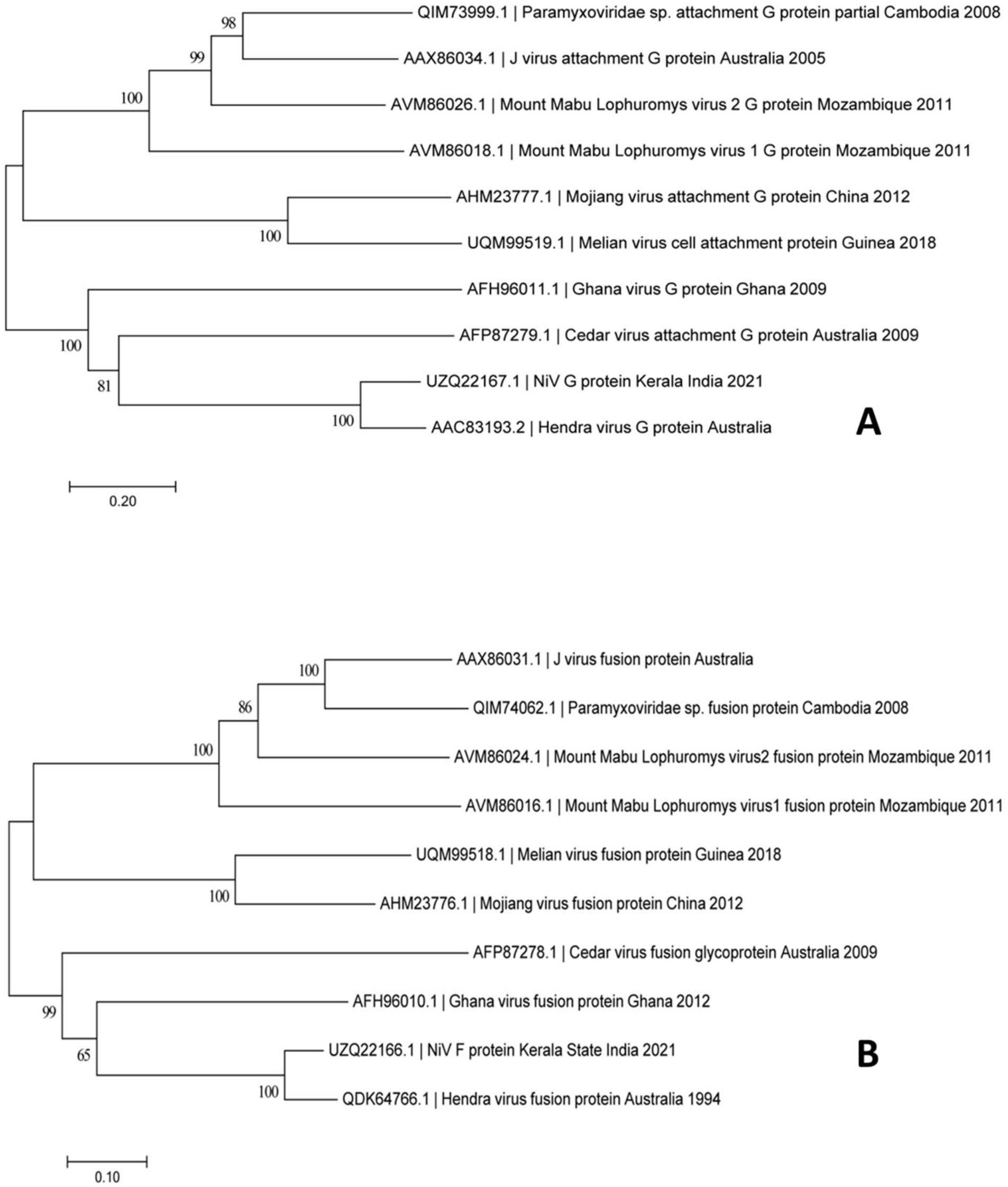
The evolutionary relationship of NiV proteins. A. The evolutionary relationship of NiVG and other viral glycoproteins using Molecular Evolutionary Genetics Analysis 7. B. The evolutionary relationship of NiVF and other viral fusion proteins using Molecular Evolutionary Genetics Analysis 7.

Figure 2B represents the evolutionary relationship of NiVF with other viral fusion proteins. The most appropriate tree is displayed with a sum of branch lengths at 3.22, and using UZQ22166.1, a recently reported NiVF protein sequence, taken here from Figure S2 [26]. The viral fusion glycoproteins included in this chart are: The NiVF protein sequence along with the fusion protein sequences of Cedar virus, Ghana virus, Hendra virus, J virus, Melian virus, Mojiang virus, Mount Mabu Lophuromys virus1 and virus2 and Paramyxoviridae sp. [8, 26–34].

The phylogenetic analysis indicated that the NiVG and NiVF sequence were in the same sub cluster that contained the Hendra virus. The CedPVG and -F species have lower sequence identities compared to those of NiVG/HeVG and NiVF/HeVF [8], and this is reflected in the phylogenetic charts in Figures 2 and 3. The evolutionary relationship of the 17 NiVF sequences observed in different countries are displayed in SI Figure S2. This phylogenetic analysis indicates the recent Indian NiVG and NiVF protein sequences form a separate sub cluster with the one linked to Bangladesh [35].

**Figure 3.**
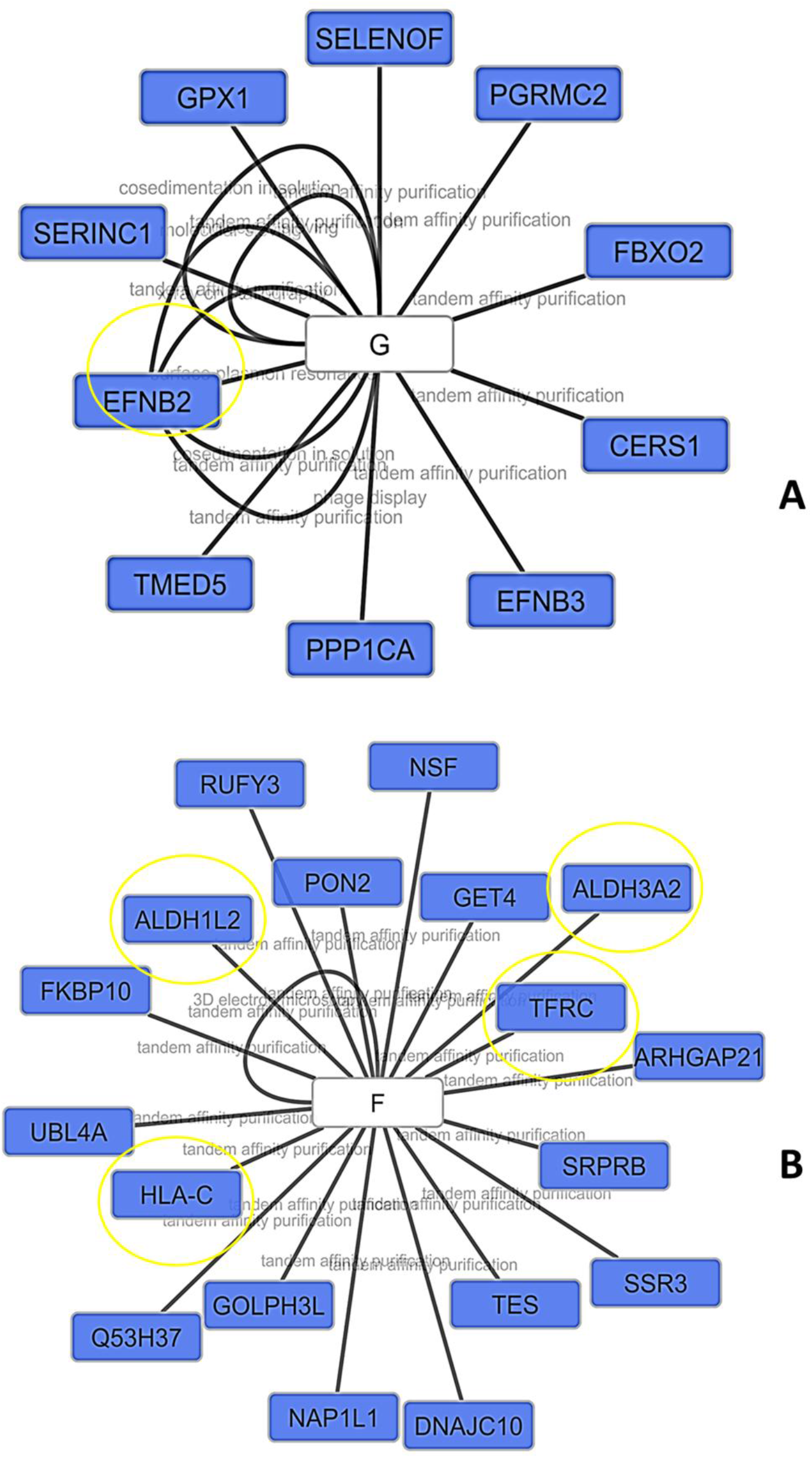
The protein-protein interactions in A. NiVG (Q9IH62) and B. NiVF (Q9IH63). These maps are based on IMEX interaction network.

The protein-protein interaction network of NiVG is demonstrated in Figure 3A, where the interaction of the glycoprotein NiVG with EFNB2, EFNB3 and other receptors are displayed using IMEx interaction network. These results have been generated using the PSICQUIC web service [36, 37], in combination with cytoscape [24]. From this network, the interactions between NiVG and EFNB2 have been identified through multiple methods, including surface plasmon resonance (SPR), X ray crystallization, tandem affinity purification, co-sedimentation procedure and phage display. Within the PPI network, the nodes represent the proteins and the edges denote interactions between the proteins. Here the number of nodes and edges are 11 and 16 respectively. Virus/host PPI network obtained using the VirHostNet2.0 knowledgebase system is included in this figure [38]. Here the protein-protein confidence score virhostnet miscore between Q9IH62 and EFNB2 is 0.327. The virhostnet represent the virus/host molecular interactions data, and the miscore represents the confidence score. The complete interactome of NiVG (Q9IH62) is displayed in Figure S3. The full network shown in Figure S3 has 7626 nodes and 28814 edges.

The protein-protein interaction network of NiVF is depicted in Figure 3B, where the interactions between NiVF and several other proteins are identified; the latter proteins include the ubiquitin-dependent protein (UBL4A), vesicle-fusing ATPase (NSF), Cgi-20 (GET4), Testin 1 (TES), etc. This network has also been generated by using cytoscape, where the numbers of both the nodes and the edges are 19. These data are also based on the IMEx interaction network and obtained from the PSICQUIC web service. In this case, the potential NiVF therapeutic targets could include the human leukocyte antigen-C (HLA-C), the aldehyde dehydrogenase 3 family member A2 (ALDH3A2), and the transferrin receptor (TFRC). The drugbank accession number for the HLA-C, ALDH1L2/ALDH3A2 and TFRC are DB02740, DB00116/DB00157 and DB05260/ DB06784/ DB01592, respectively. From this network, the interactions between NiVF and HLA-C are identified by tandem affinity purification. The interactions between NiVF and ALDH1L2/ ALDH3A2 are indicated by 3D electron microscopy and/or tandem affinity purification. The interactions between NiVF and TFRC are also characterized by tandem affinity purification. While NiVF is a target for vaccines and therapeutics development, no readily available complete interactome networks for Q9IH63 could be found at the time of this investigation.

The ectodomain of EFNB2 receptor protein consists of eight β-stranded barrels. The EFNB2 G-H loop is a conserved loop region that is critical for binding the henipavirus (Hendra virus and Nipah virus) G-proteins [6, 39–42]. The flexible G-H loop is located between βG and βH, and projected above the surface of the receptor. The viral G-proteins undergo crucial conformational changes after receptor binding. Bowden et al have suggested that in the ligand bound form, the G-H loop of EFNB2 receptor makes complementary binding with NiVG and that a stable NiVG-EFNB2 complex is formed via the “induced-fit” mechanism. [39].

The G-H loop residues Glu118-Leu127 can be considered as targets for antibody neutralization. The RMSF data (presented later in this report) demonstrate that these G-H loop residues are quite stable in the NiVG-EFNB2 complex. According to the recent literature, several neutralization vaccines bind to EFN G-H and thus, interrupt the interactions between the receptor and the Henipavirus glycoprotein. Therefore, it is essential to identify the interfacial residues within these proteins that could be utilized to neutralize the ligand receptor interactions. The interfacial residues in NiVG and NiVF are considered in Tables 1 and 2. The NiVG and NiVF non-bond monitors in 2VSM and 6T3F are tabulated in SI Tables S1-S4. The unfavorable non-bond monitors of these two ligands are also tabulated there. 2D structural analyses of the crucial G-H loop residues 2VSM: B are displayed in SI Table S5.

**Table 1.**
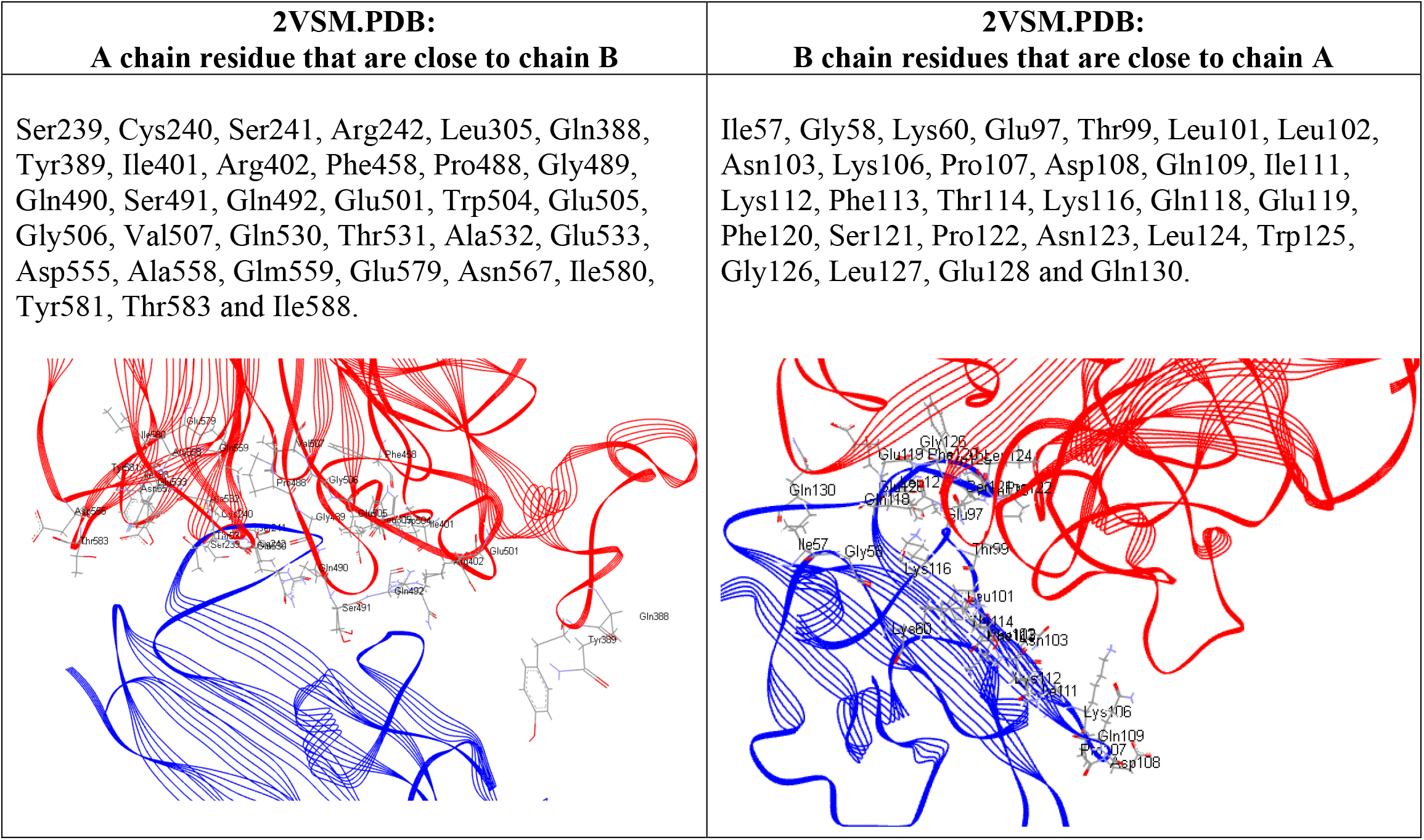
Protein-protein interfacial residues in 2VSM.PDB. The truncated form of 2VSM, that is used in the MD simulation is analyzed here. Chain A (red) is the NiV glycoprotein and chain B (blue) represents the Ephrin-B2 receptor. The interfacial residues are represented in stick mode and atoms are colored by elements.

| **2VSM.PDB:**<br>**A chain residue that are close to chain B** | **2VSM.PDB:**<br>**B chain residues that are close to chain A** |
| --- | --- |
| Ser239, Cys240, Ser241, Arg242, Leu305, Gln388, Tyr389, Ile401, Arg402, Phe458, Pro488, Gly489, Gln490, Ser491, Gln492, Glu501, Trp504, Glu505, Gly506, Val507, Gln530, Thr531, Ala532, Glu533, Asp555, Ala558, Glm559, Glu579, Asn567, Ile580, Tyr581, Thr583 and Ile588. | Ile57, Gly58, Lys60, Glu97, Thr99, Leu101, Leu102, Asn103, Lys106, Pro107, Asp108, Gln109, Ile111, Lys112, Phe113, Thr114, Lys116, Gln118, Glu119, Phe120, Ser121, Pro122, Asn123, Leu124, Trp125, Gly126, Leu127, Glu128 and Gln130. |

**Table II.**
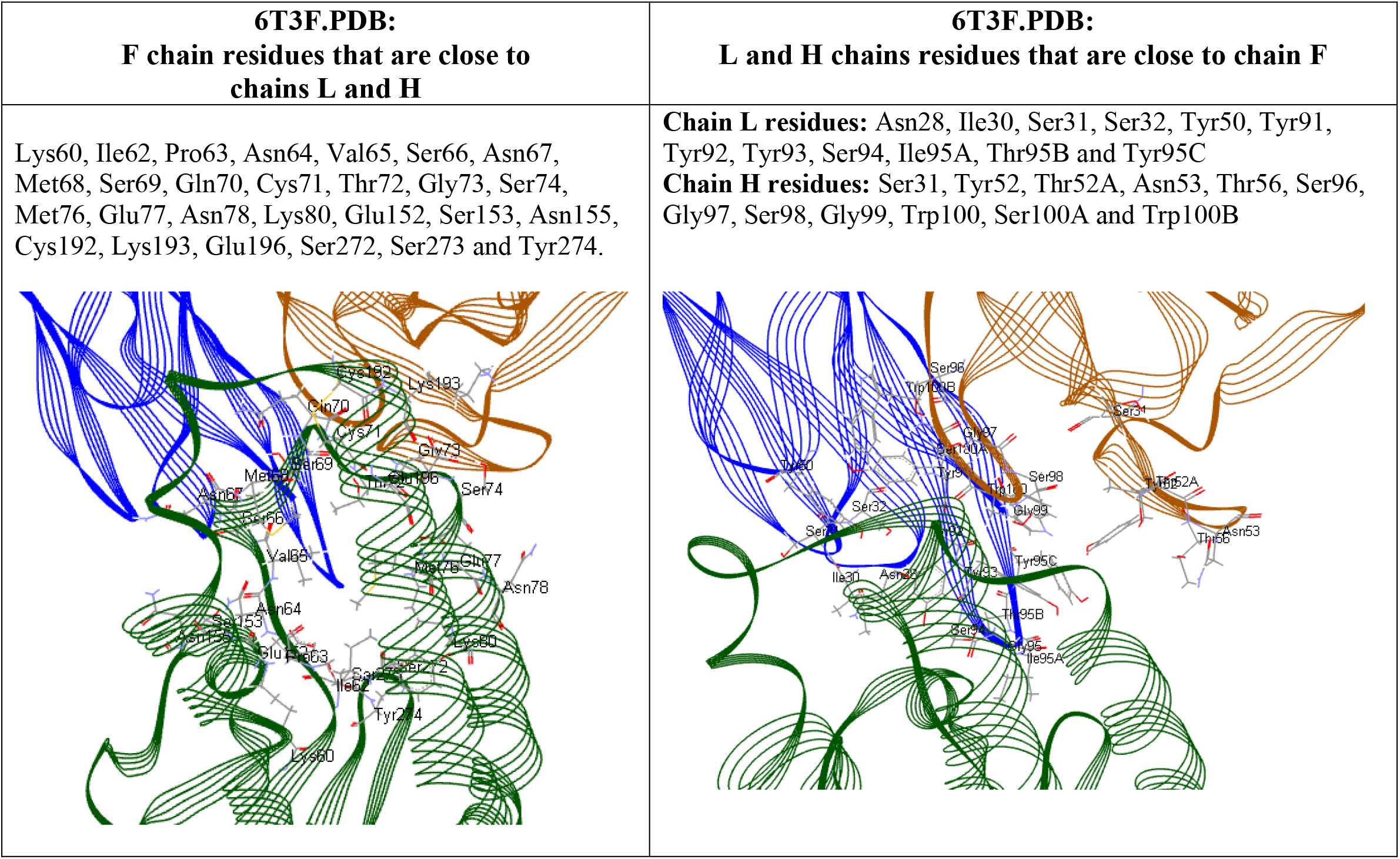
Interfacial residues in 6T3F.PDB. Chain F (green) is the NiV fusion protein and chains L (blue) and H (brown) represent the light and heavy chains of Fab66 receptor. The truncated form of 6T3F, used in the MD simulation is analyzed here. The interfacial residues are represented in stick mode and atoms are colored by elements.

| **6T3F.PDB:**<br>**F chain residues that are close to chains L and H** | **6T3F.PDB:**<br>**L and H chains residues that are close to chain F** |
| --- | --- |
| Lys60, Ile62, Pro63, Asn64, Val65, Ser66, Asn67, Met68, Ser69, Gln70, Cys71, Thr72, Gly73, Ser74, Met76, Glu77, Asn78, Lys80, Glu152, Ser153, Asn155, Cys192, Lys193, Glu196, Ser272, Ser273 and Tyr274. | **Chain L residues:** Asn28, Ile30, Ser31, Ser32, Tyr50, Tyr91, Tyr92, Tyr93, Ser94, Ile95A, Thr95B and Tyr95C<br>**Chain H residues:** Ser31, Tyr52, Thr52A, Asn53, Thr56, Ser96, Gly97, Ser98, Gly99, Trp100, Ser100A and Trp100B |

The RMSD value represents a measure of the protein’s structural deviation from its starting structure (at 0 ns) during the MD simulation; lower RMSD values are attributed to temporally stable and steady state characteristics of a protein. Figure 4 shows the all-atom RMSD plots for 2VSM.PDB and 6T3F.PDB, which indicate adequate temporal stability of these proteins. However, the protein complex of NiVG with EFNB2 receptor (2VSM) exhibits a relatively lower RMSD an average value of 3.19 Å ±0.28 Å, with maximum and minimum RMSDs at 3.85 Å and 1.04 Å, respectively) compared to that of the NiVF protein with Fab receptor, 6T3F.PDB. For 6T3F (which yields an overall average RMSD value of 4.37 Å ±0.37Å, with maximum and minimum RMSDs being 5.81Å and 1.30Å, respectively).

**Figure 4.**
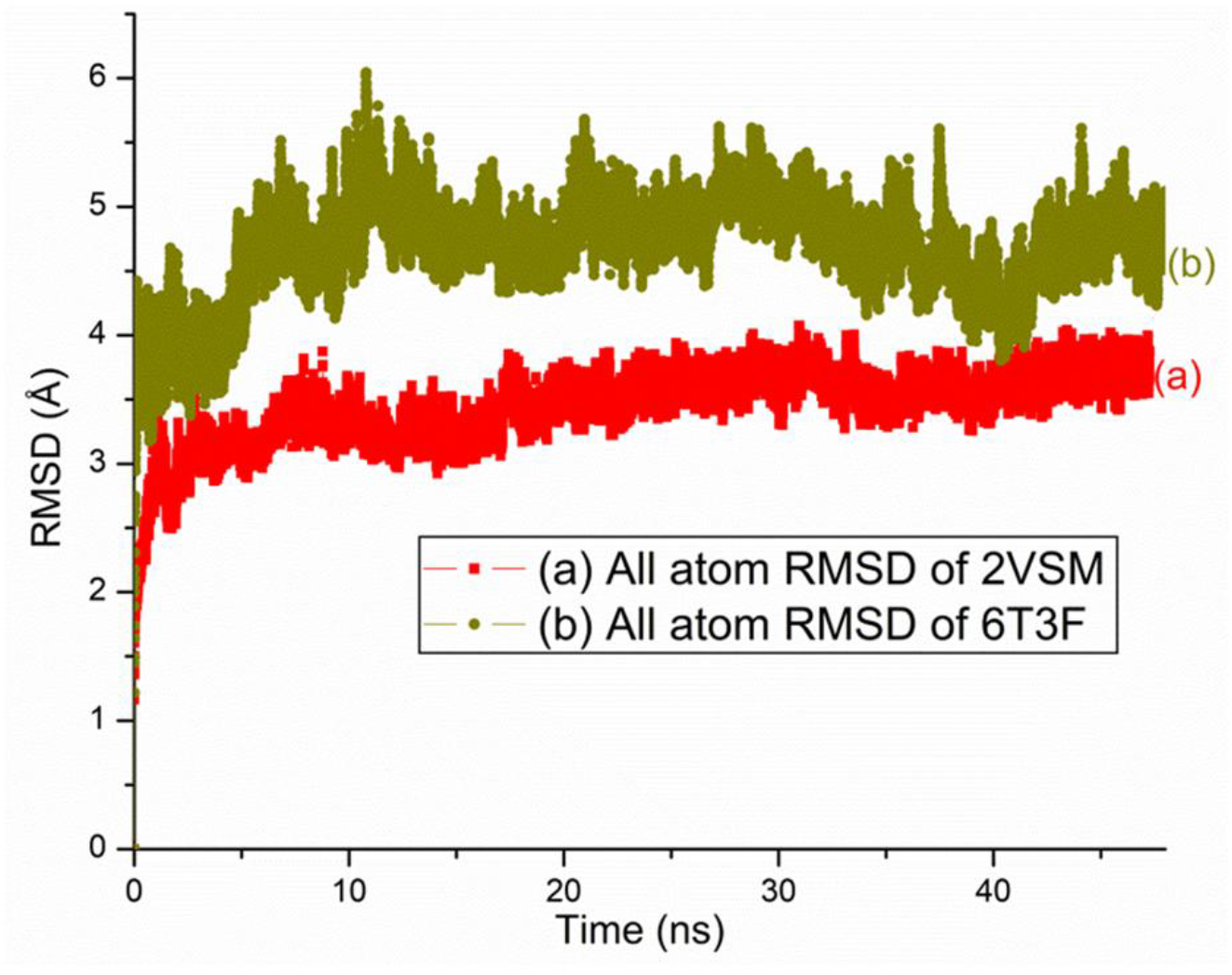
All atom RMSD plots of 2VSM (a; red); and 6T3F (b; olive green).

It is possible that residues 101-116, 161-168, 214-226, and 248-252 form “turn and loop” like structures in 6T3F in addition to several other short turns and long loops between helices and strands that exist within the 6T3F. Few residues of the F chain also exhibit higher fluctuations and movements; this likely happens due to their presence at surface regions, where the strength of protein-protein interactions are weak, and hence the overall structural stability of the proteins might be limited. The ligand-receptor enabled fit mechanism in 2VSM.PDB makes this protein relatively more stable, and this is reflected in the observed RMSD values. Additionally, 2VSM has several more beta stands compare to the those of the truncated 6T3F structure, and this makes the 2VSM comparatively more stable. RMSD plots of these two proteins obtained in an explicit solvent system are shown in the SI Figure S4. This exploratory explicit RMSD values are correlated to the RMSD data in an implicit solvent model.

Figure 5 represent the RMSF plots of the above two protein chains. These plots identify the flexible regions of the protein, and the RMSF values represent the average deviation of amino acid residues (within the protein structure) from their mean positions during the simulation time. The higher fluctuations correspond to higher RMSF values, while the stable areas are characterized by lower RMSF values. In 2VSM glycoprotein, chain A is stable except for the initial few residues and the residues, 420-421 (Figure 5A). The targeted G-H loop residues for antibody neutralization within 2VSM: B appear quite stable (Figure 5B). In the truncated 6T3F: F chain (Figure 5C), several residues, in particular residues 102-110, 117-125, 215-226, 249-252, 258-261, have higher fluctuations due to their unstable locations within turn or loop regions and/or the outer surface regions. These residues are distal to the Fab66 binding site and show greater flexibility. Truncation could also play a role in dictating the protein’s flexibility. From this Figure it appears evident that the 2VSM is a comparatively more stable structure.

**Figure 5.**
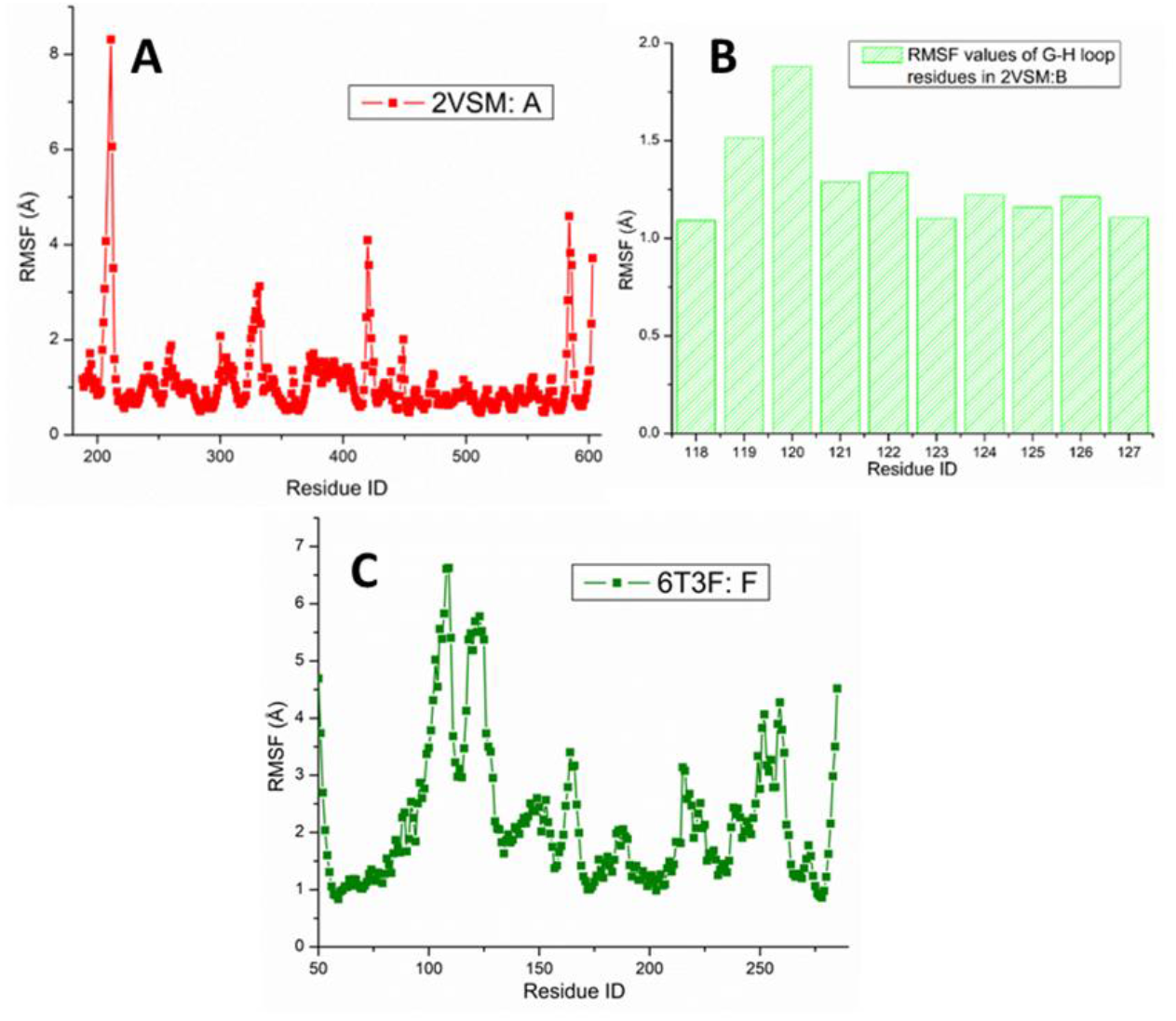
RMSF plots of A. 2VSM: chain A (red); B. some residues within G-H loop of 2VSM.PDB. and C. 6T3F: chain F (green).

## 4. Conclusions

While the number of reported NiV cases have been relatively small in recent years, and have been mostly limited to certain geographic areas, the detailed structural aspects of this virus had not been probed in details previously. While receptor ephrin-B can mediate bidirectional signaling in tumorigenesis, they independently act as an entry receptor for fatal Nipah virus infection. Using classical simulation methods, the present work demonstrates that the ephrinB2 bound NiVG is a stable protein. Within the limits of its classical formalism (lacking quantum mechanical information), the detailed analysis of the receptor bound NiV structure may help to identify the hotspots that are responsible for protein’s stability. The present results demonstrate the applicability of molecular dynamics simulations to identify the functional epitopes and immune escape mutants to identify the antiviral candidates. The structural features of the NiV reported in this work are expected to be useful in the investigative context of vaccine development strategies. The presence of multiple sequence alignment within genre Henipavirus, suggests that HeVF and NiVF have high sequence similarity and conserved sequence [43]. Additionally, the evolutionary relationship between NiVG and NiVF (Figure 2A and B) shows a close association of NiV with HeV and the CedPV within the Henipavirus genre. This phylogenetic analysis is important to detect mutational mapping and classification of future lineages.

For a long time, artificial intelligence (AI) and machine learnings (ML) based techniques have been used to detect, diagnose and treat various diseases. In recent years AI based approaches like AlphaFold and RoseTTAFold have been applied to predict proteins’ 3D structures when appropriate experimental NMR, X-ray or cryo-electron microscopy (cryo-EM) structures are unavailable [44, 45]. These AI based techniques are applied to predict structures from the evolutionary data and proteins’ sequences. The data driven AI based technologies can also identify the molecular level protein-protein interactions and proteins’ network prediction. These predictive approaches can be used to further determine viral evolutions and virus variants. Based on the protein’s interfacial interactions, discussed in this paper the AI can be applied to this study for identifying antigenic epitopes, antigen-adjuvant combination and vaccine candidates for NiV and other emerging zoonotic diseases.

Supplementary Information, Figures S1-S4; Tables S1-S5.

## Statements and Declarations

The author did not receive support from any organization for the submitted work. The author has no relevant financial or non-financial interests to disclose.

## Acknowledgments

The author acknowledges utilization of the following simulation and visualization software packages: 1) NAMD and 2) VMD: NAMD and VMD, developed by the Theoretical and Computational Biophysics Group in the Beckman Institute for Advanced Science and Technology at the University of Illinois, Urbana-Champaign. 3) Discovery Studio Visualizer: Dassault Systèmes BIOVIA, Discovery Studio Modeling Environment, San Diego, CA: Dassault Systèmes (2015).

