## Supplementary Information for "Structural Aspects of Nipah Virus Glycoproteins using Predictive Protein Modeling"

for

### Structural Aspects of Nipah Virus Glycoproteins

Urmi Roy*

*Department of Chemistry & Biochemistry
Clarkson University
8 Clarkson Avenue, Potsdam, NY 13699, United States

ORCiD ID: https://orcid.org/0000-0003-0928-0574

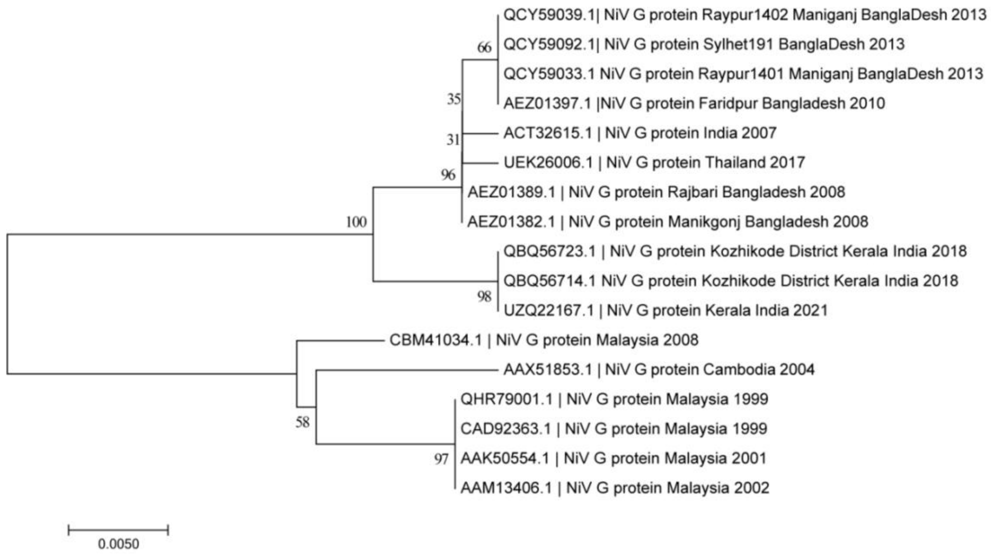


Fig. S1. The evolutionary relationship of 17 Nipah virus glycoproteins (G proteins) from different countries using Molecular Evolutionary Genetics Analysis 7 (MEGA 7) [1]. Here the evolutionary tree is presented in the traditional way. The total positions in the final dataset are 562. Sum of branch length is 0.071. The Bootstrap method was used for the test of phylogeny. This evolutionary history was inferred using the Neighbor-Joining method [2]. Any gaps or missing data were deleted. The number of bootstrap replications was 200 [3]. The Poisson model was used to calculate the evolutionary distances [4].

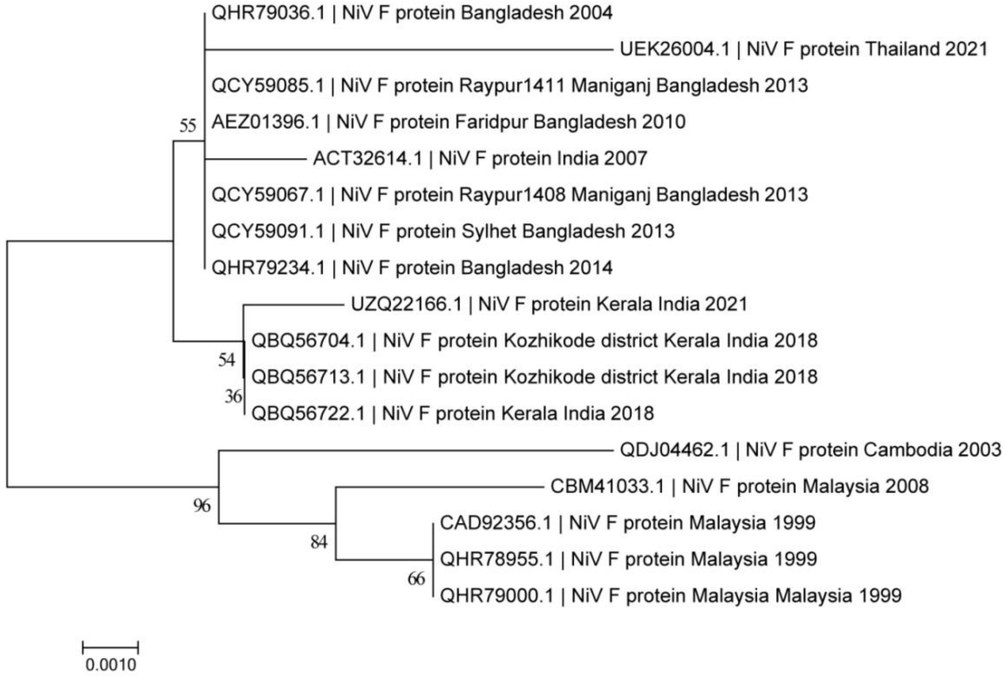


Fig. S2. The evolutionary relationship of 17 Nipah virus Fusion glycoproteins from different countries using Molecular Evolutionary Genetics Analysis 7 (MEGA 7) [1]. The total positions in the final dataset are 544. Sum of branch length is 0.035. The Bootstrap method was used for the test of phylogeny. This evolutionary history was inferred using the Neighbor-Joining method [2]. Any gaps or missing data were deleted. The number of bootstrap replications was 200 [3]. The Poisson model was used to calculate the evolutionary distances [4]. Figure S1-S2 represent the evolutionary relationship of the Nipah virus glycoprotein and fusion protein from different countries including Bangladesh [5-7], Cambodia [8-9], India [10-12], Malaysia [13-17] and Thailand [18].

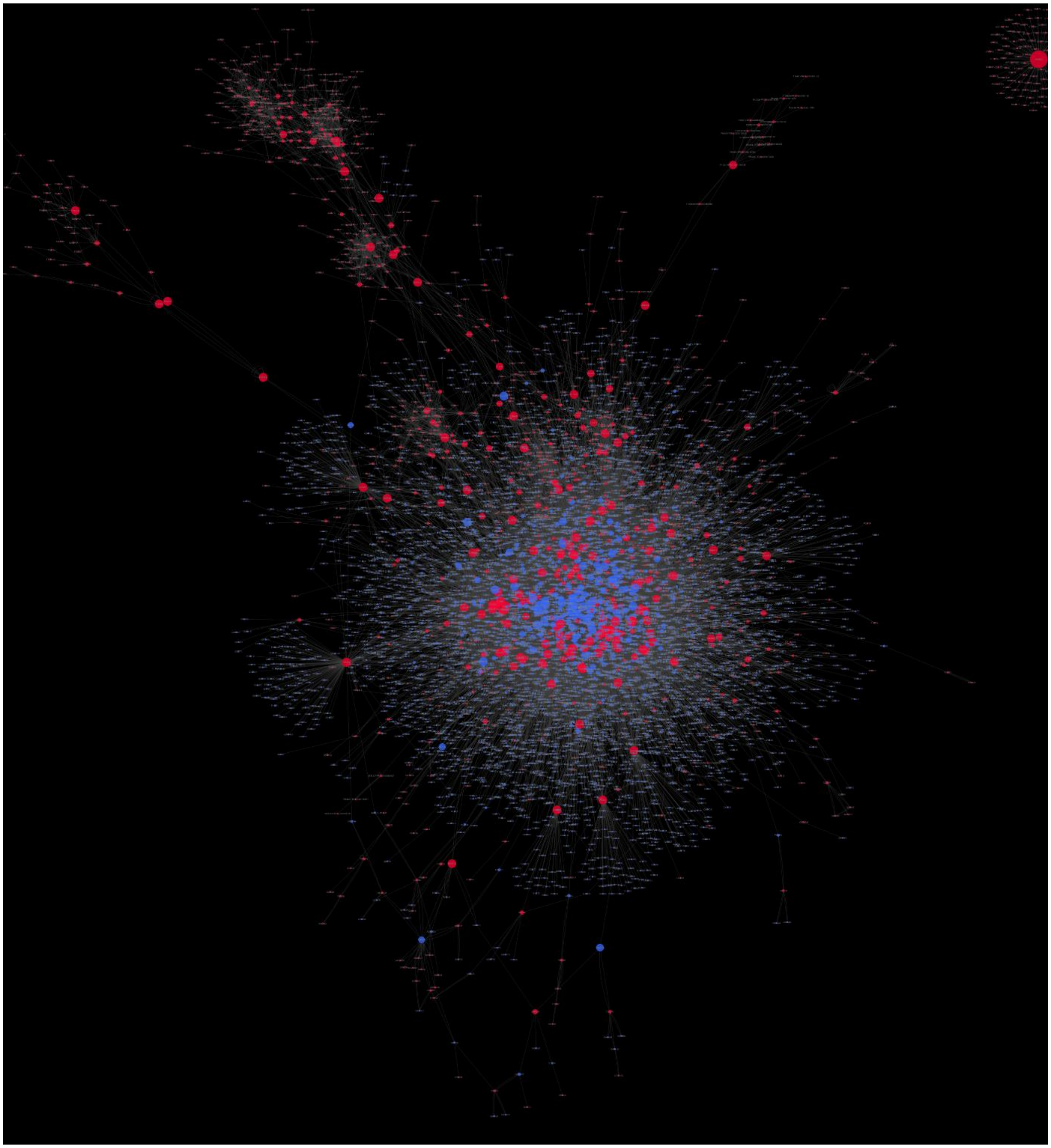

Figure S3. The interactome network of NiVG. Here the edge betweenness is 28084 and shared interactions are 25017. This analysis was done by Cytoscape [19]

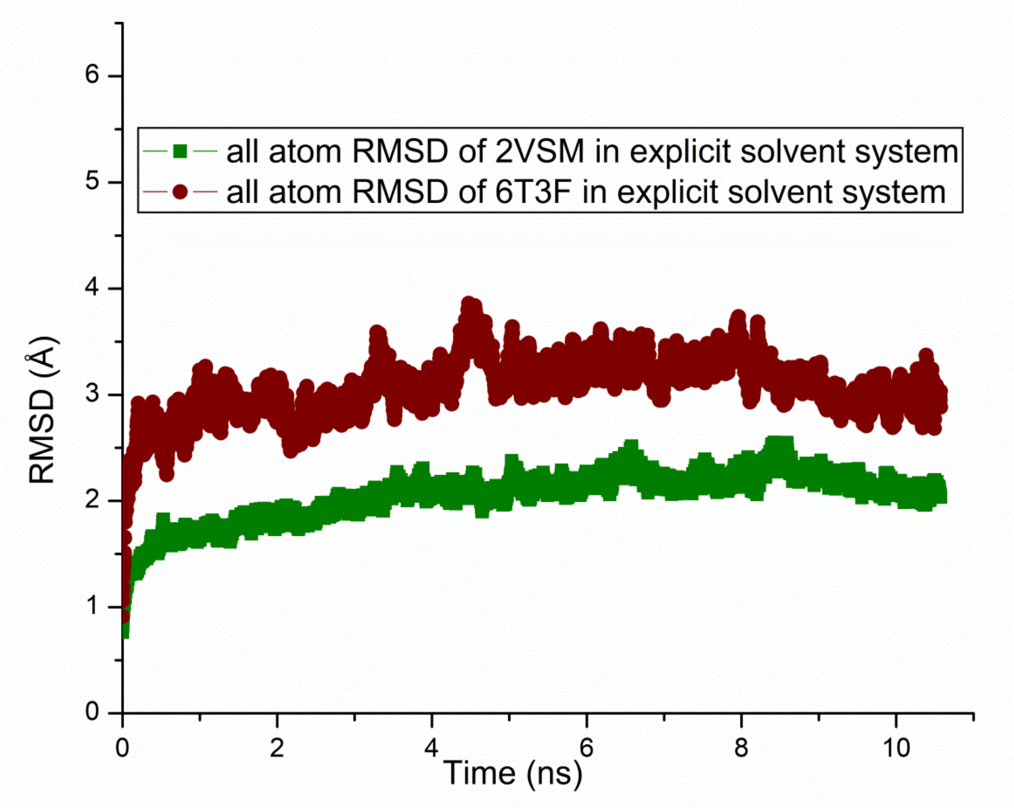


Figure S4. The RMSD plots of 2VSM and 6T3F.

The MD simulation of these two systems was performed using NAMD, VMD and QwikMD [20-22]. All simulations were carried out in an explicit solvent system using NpT ensemble and using the TIP3 water model [23]. These two systems were solvated in a cubic water box with 15 Å buffer. They were neutralized using sodium and chloride ions where the salt concentration was 0.15 mol/L. The CHARMM36 force field was applied in both simulations [24]. The system energy was minimized in 2000 steps. After the minimization steps, the annealing process was applied for 0.24 ns and the temperature of the system was ramped up from 60 K to 300 K. The equilibration steps were for 0.04 ns and then the final MD simulation was run for 10 ns.

**Table S1.** Ligand non-bond monitor in 2VSM

| 2VSM: non bond interaction | Distance | Category | Types | from Chemistry | to Chemistry |
|---|---|---|---|---|---|
| B:LYS60:HZ3 - A:GLU533:OE1 | 1.93947 | Hydrogen Bond;Electrostatic | Salt Bridge;Attractive Charge | H-Donor;Positive | H-Acceptor;Negative |
| B:LYS106:HZ1 - A:GLU501:OE2 | 1.75478 | Hydrogen Bond;Electrostatic | Salt Bridge;Attractive Charge | H-Donor;Positive | H-Acceptor;Negative |

| | | | | | |
|---|---|---|---|---|---|
| B:LYS116:HZ3 - A:GLU533:OE2 | 1.92533 | Hydrogen Bond;Electrostatic | Salt Bridge;Attractive Charge | H-Donor;Positive | H-Acceptor;Negative |
| B:LYS116:NZ - A:ASP555:OD2 | 5.54879 | Electrostatic | Attractive Charge | Positive | Negative |
| A:CYS240:HN - B:GLU119:OE2 | 1.72848 | Hydrogen Bond | Conventional Hydrogen Bond | H-Donor | H-Acceptor |
| A:CYS240:HN - B:GLU128:OE1 | 3.05815 | Hydrogen Bond | Conventional Hydrogen Bond | H-Donor | H-Acceptor |
| A:SER241:HN - B:GLU128:OE2 | 2.00286 | Hydrogen Bond | Conventional Hydrogen Bond | H-Donor | H-Acceptor |
| A:ARG242:HH21 - B:TRP125:O | 2.12932 | Hydrogen Bond | Conventional Hydrogen Bond | H-Donor | H-Acceptor |
| A:GLN388:HE22 - B:ASP108:OD2 | 2.68141 | Hydrogen Bond | Conventional Hydrogen Bond | H-Donor | H-Acceptor |
| A:ARG402:HE:B - B:GLU97:O | 1.92098 | Hydrogen Bond | Conventional Hydrogen Bond | H-Donor | H-Acceptor |
| A:ARG402:HH21:B - B:GLU97:O | 2.51411 | Hydrogen Bond | Conventional Hydrogen Bond | H-Donor | H-Acceptor |
| A:SER491:HG:B - B:LEU101:O | 2.39037 | Hydrogen Bond | Conventional Hydrogen Bond | H-Donor | H-Acceptor |
| A:GLN492:HE22 - B:ASN103:OD1 | 2.23171 | Hydrogen Bond | Conventional Hydrogen Bond | H-Donor | H-Acceptor |
| A:GLY506:HN - B:PRO122:O | 1.89097 | Hydrogen Bond | Conventional Hydrogen Bond | H-Donor | H-Acceptor |
| A:GLN530:HE22 - B:LYS112:O | 1.98504 | Hydrogen Bond | Conventional Hydrogen Bond | H-Donor | H-Acceptor |
| B:THR114:HN - A:GLN530:OE1 | 1.93673 | Hydrogen Bond | Conventional Hydrogen Bond | H-Donor | H-Acceptor |
| B:LYS116:HZ1 - A:ASP555:O | 2.43272 | Hydrogen Bond | Conventional Hydrogen Bond | H-Donor | H-Acceptor |
| B:LYS116:HZ2 - A:ASN557:OD1 | 1.77832 | Hydrogen Bond | Conventional Hydrogen Bond | H-Donor | H-Acceptor |
| B:GLN118:HE22 - A:ALA532:O | 1.96124 | Hydrogen Bond | Conventional Hydrogen Bond | H-Donor | H-Acceptor |
| B:GLU119:HN - A:TYR581:OH | 1.8949 | Hydrogen Bond | Conventional Hydrogen Bond | H-Donor | H-Acceptor |

| | | | | | |
|---|---|---|---|---|---|
| B:SER121:HN - A:GLN559:OE1 | 2.00425 | Hydrogen Bond | Conventional Hydrogen Bond | H-Donor | H-Acceptor |
| B:ASN123:HD21 - A:GLN490:OE1 | 1.96527 | Hydrogen Bond | Conventional Hydrogen Bond | H-Donor | H-Acceptor |
| A:SER239:HA - B:GLU119:OE2 | 2.77701 | Hydrogen Bond | Carbon Hydrogen Bond | H-Donor | H-Acceptor |
| A:SER239:HB1 - B:GLU119:OE2 | 2.92813 | Hydrogen Bond | Carbon Hydrogen Bond | H-Donor | H-Acceptor |
| A:SER239:HB1 - B:GLU128:OE2 | 2.32143 | Hydrogen Bond | Carbon Hydrogen Bond | H-Donor | H-Acceptor |
| A:SER241:HB1 - B:GLY126:O | 2.89636 | Hydrogen Bond | Carbon Hydrogen Bond | H-Donor | H-Acceptor |
| A:SER241:HB1 - B:GLU128:OE2 | 2.15657 | Hydrogen Bond | Carbon Hydrogen Bond | H-Donor | H-Acceptor |
| A:SER241:HB2 - B:GLY126:O | 2.74683 | Hydrogen Bond | Carbon Hydrogen Bond | H-Donor | H-Acceptor |
| A:SER491:HA:B - B:LYS112:O | 2.94971 | Hydrogen Bond | Carbon Hydrogen Bond | H-Donor | H-Acceptor |
| A:SER491:HB1:B - B:LYS112:O | 2.43778 | Hydrogen Bond | Carbon Hydrogen Bond | H-Donor | H-Acceptor |
| A:SER491:HB2:B - B:LEU101:O | 2.9488 | Hydrogen Bond | Carbon Hydrogen Bond | H-Donor | H-Acceptor |
| A:SER491:HA - B:LYS112:O | 2.86276 | Hydrogen Bond | Carbon Hydrogen Bond | H-Donor | H-Acceptor |
| A:SER491:HB1 - B:LEU101:O | 2.77501 | Hydrogen Bond | Carbon Hydrogen Bond | H-Donor | H-Acceptor |
| A:SER491:HB2 - B:LEU101:O | 2.65631 | Hydrogen Bond | Carbon Hydrogen Bond | H-Donor | H-Acceptor |
| A:GLU505:HA - B:PRO122:O | 2.93327 | Hydrogen Bond | Carbon Hydrogen Bond | H-Donor | H-Acceptor |
| B:LYS60:HE2 - A:GLU533:OE1 | 2.70241 | Hydrogen Bond | Carbon Hydrogen Bond | H-Donor | H-Acceptor |
| B:PHE113:HA - A:GLN530:OE1 | 2.41693 | Hydrogen Bond | Carbon Hydrogen Bond | H-Donor | H-Acceptor |
| B:LYS116:HE1 - A:ASN557:OD1 | 2.78569 | Hydrogen Bond | Carbon Hydrogen Bond | H-Donor | H-Acceptor |

| | | | | | |
|---|---|---|---|---|---|
| B:PHE120:HA - A:GLN559:OE1 | 2.52772 | Hydrogen Bond | Carbon Hydrogen Bond | H-Donor | H-Acceptor |
| B:PRO122:HA - A:GLY506:O | 2.67339 | Hydrogen Bond | Carbon Hydrogen Bond | H-Donor | H-Acceptor |
| A:ARG402:NH1 - B:TRP125 | 3.63704 | Electrostatic | Pi-Cation | Positive | Pi-Orbitals |
| A:TRP504 - B:TRP125 | 4.91734 | Hydrophobic | Pi-Pi T-shaped | Pi-Orbitals | Pi-Orbitals |
| A:TYR581 - B:PHE120 | 5.18914 | Hydrophobic | Pi-Pi T-shaped | Pi-Orbitals | Pi-Orbitals |
| B:TRP125 - A:TRP504 | 5.16202 | Hydrophobic | Pi-Pi T-shaped | Pi-Orbitals | Pi-Orbitals |
| A:PRO488 - B:PRO122 | 5.04236 | Hydrophobic | Alkyl | Alkyl | Alkyl |
| A:TYR389 - B:LYS106 | 3.87551 | Hydrophobic | Pi-Alkyl | Pi-Orbitals | Alkyl |
| A:PHE458 - B:LEU124 | 4.56962 | Hydrophobic | Pi-Alkyl | Pi-Orbitals | Alkyl |
| A:TRP504 - B:LEU124 | 4.42722 | Hydrophobic | Pi-Alkyl | Pi-Orbitals | Alkyl |
| A:TRP504 - B:LEU124 | 4.22236 | Hydrophobic | Pi-Alkyl | Pi-Orbitals | Alkyl |
| B:TRP125 - A:ILE401 | 5.08014 | Hydrophobic | Pi-Alkyl | Pi-Orbitals | Alkyl |

**Table S2.** Unfavorable non-bond monitor in 2VSM ligand

| *2VSM: unfavourable non bond* | *Distance* | *Category* | *Types* | *from Chemistry* | *to Chemistry* |
|---|---|---|---|---|---|
| *A:SER491:OG - B:LYS112:O* | *2.88876* | *Unfavorable* | *Unfavorable Acceptor-Acceptor* | *H-Acceptor* | *H-Acceptor* |

**Table S3.** Ligand non-bond monitor in 6T3F

| **6T3F: non bond interaction** | **Distance** | **Name** | **Types** | **From Chemistry** | **To Chemistry** |
|---|---|---|---|---|---|
| F:VAL65:HN - L: SER94:O | 2.16621 | Hydrogen Bond | Conventional Hydrogen Bond | H-Donor | H-Acceptor |
| F:SER69:HG - L:TYR50:OH | 2.71456 | Hydrogen Bond | Conventional Hydrogen Bond | H-Donor | H-Acceptor |
| F:GLY73:HN - H:SER98:OG | 2.88496 | Hydrogen Bond | Conventional Hydrogen Bond | H-Donor | H-Acceptor |
| F:LYS80:HZ2 - L:GLY95:O | 2.08184 | Hydrogen Bond | Conventional Hydrogen Bond | H-Donor | H-Acceptor |

| | | | | | |
|---|---|---|---|---|---|
| L:TYR50:HH - F:SER66:O | 2.8748 | Hydrogen Bond | Conventional Hydrogen Bond | H-Donor | H-Acceptor |
| L:TYR95C:HH - F:GLU77:OE2 | 2.09196 | Hydrogen Bond | Conventional Hydrogen Bond | H-Donor | H-Acceptor |
| H:THR52A:HG1 - F:SER74:OG | 1.90095 | Hydrogen Bond | Conventional Hydrogen Bond | H-Donor | H-Acceptor |
| H:SER98:HN - F:GLN70:O | 2.40343 | Hydrogen Bond | Conventional Hydrogen Bond | H-Donor | H-Acceptor |
| H:SER98:HG - F:GLN70:O | 2.5759 | Hydrogen Bond | Conventional Hydrogen Bond | H-Donor | H-Acceptor |
| H:GLY99:HN - F:GLN70:O | 1.62535 | Hydrogen Bond | Conventional Hydrogen Bond | H-Donor | H-Acceptor |
| F:ASN64:HA - L:SER94:O | 2.77495 | Hydrogen Bond | Carbon Hydrogen Bond | H-Donor | H-Acceptor |
| F:SER66:HA - L:TYR93:O | 2.59908 | Hydrogen Bond | Carbon Hydrogen Bond | H-Donor | H-Acceptor |
| F:SER69:HB1 - H:TRP100:O | 2.80682 | Hydrogen Bond | Carbon Hydrogen Bond | H-Donor | H-Acceptor |
| F:GLN70:HA - H:GLY99:O | 2.72538 | Hydrogen Bond | Carbon Hydrogen Bond | H-Donor | H-Acceptor |
| F:GLN70:HA - H:TRP100:O | 2.58701 | Hydrogen Bond | Carbon Hydrogen Bond | H-Donor | H-Acceptor |
| F:LYS80:HE1 - L:GLY95:O | 2.90219 | Hydrogen Bond | Carbon Hydrogen Bond | H-Donor | H-Acceptor |
| L:SER94:HA - F:ASN64:OD1 | 2.40998 | Hydrogen Bond | Carbon Hydrogen Bond | H-Donor | H-Acceptor |
| H:GLY97:HA2 - F:GLN70:O | 2.50417 | Hydrogen Bond | Carbon Hydrogen Bond | H-Donor | H-Acceptor |
| H:GLY99:HA2 - F:SER69:O | 2.41558 | Hydrogen Bond | Carbon Hydrogen Bond | H-Donor | H-Acceptor |
| F:ILE62 - L:ILE95A | 4.80116 | Hydrophobic | Alkyl | Alkyl | Alkyl |

**Table S4.** Unfavorable non-bond monitor in 6T3F ligand.
*The interactions and interaction analyses in Table1-6 were generated using Discovery Studio Visualizer [20].*

| Unfavorable non bond in 6T3F | Distance | Category | Types | From Chemistry | To Chemistry |
|---|---|---|---|---|---|
| F:ASN64:HD22 - L:TYR92:HH | 2.11877 | Unfavorable | Unfavorable Donor-Donor | H-Donor | H-Donor |
| F:GLU196:OE1 - H:SER98:OG | 2.96795 | Unfavorable | Unfavorable Acceptor-Acceptor | H-Acceptor | H-Acceptor |

**Table S5 [Shown on SEM pages 9-11].** Detailed structural analyses of most of the residues in G-H loop in 2VSM: B. 2D interactions of residues Glu119 and Glu128 are not shown. Since these residues have fractional formal charges and/or partial bonds.

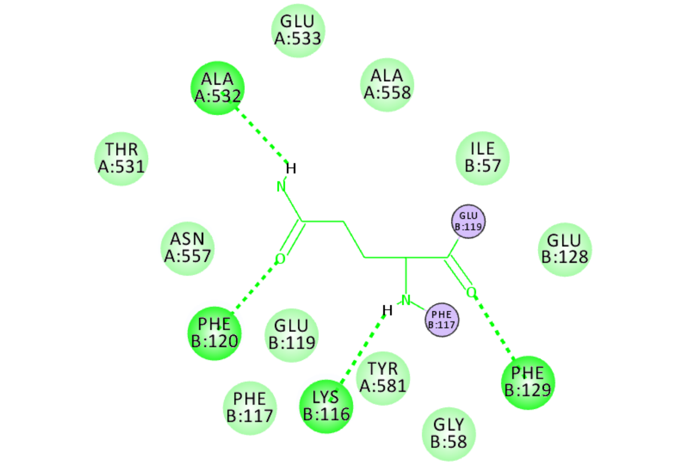


**B: Gln118**

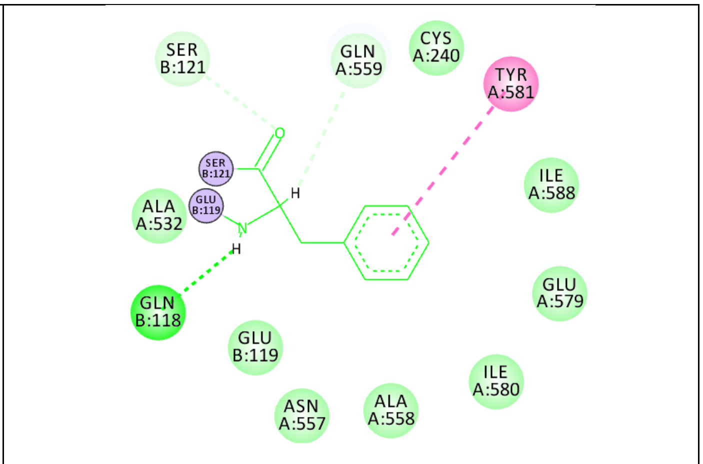


**B: Phe120**

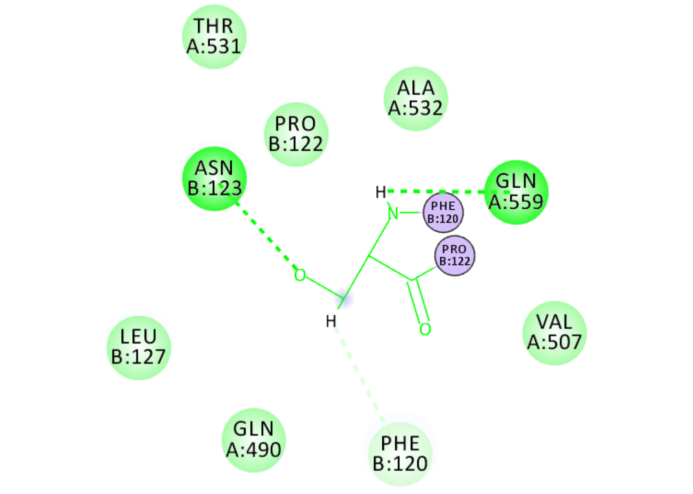


**B: Ser121**

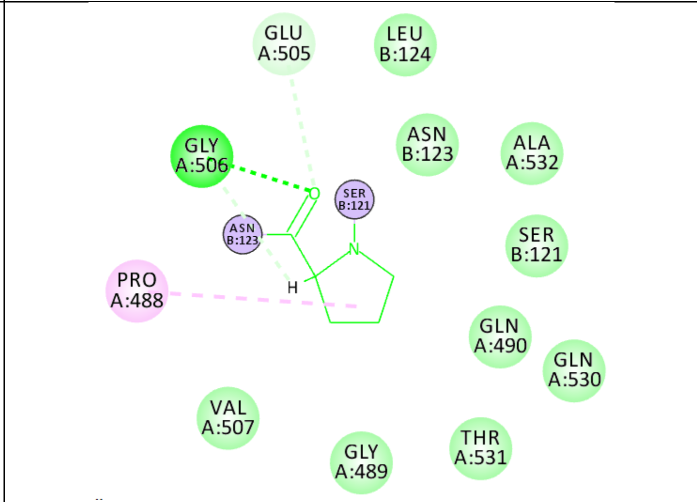


**B: Pro122**

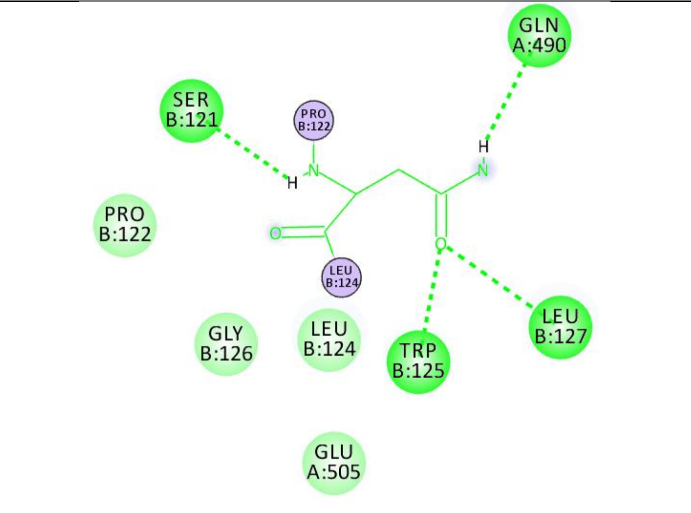


B: Asm123

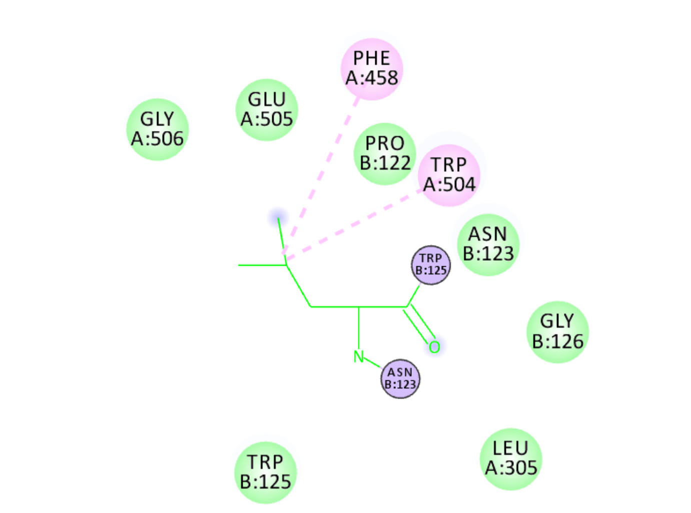


B: Leu124

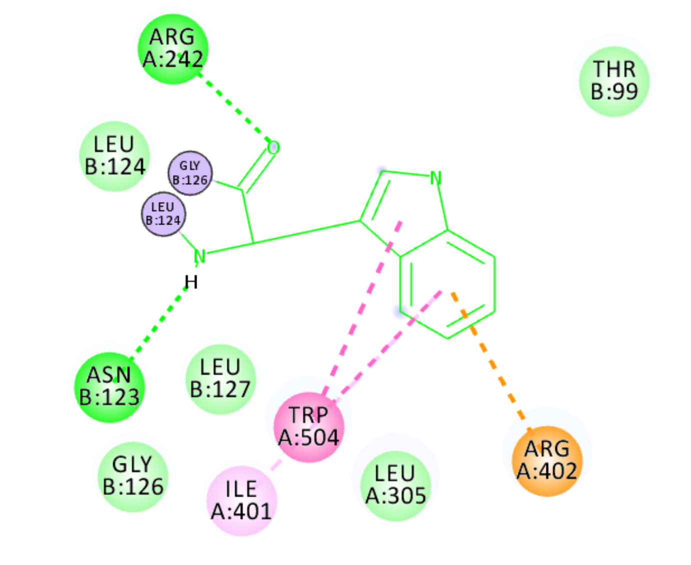


B: Trp125

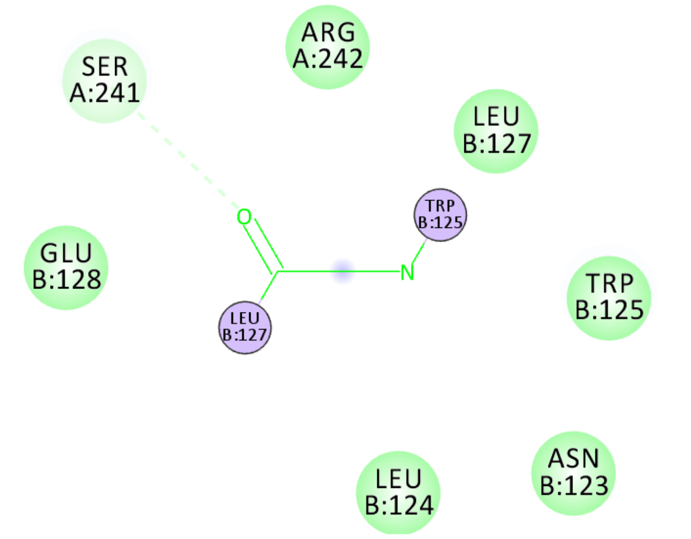


B: Gly126

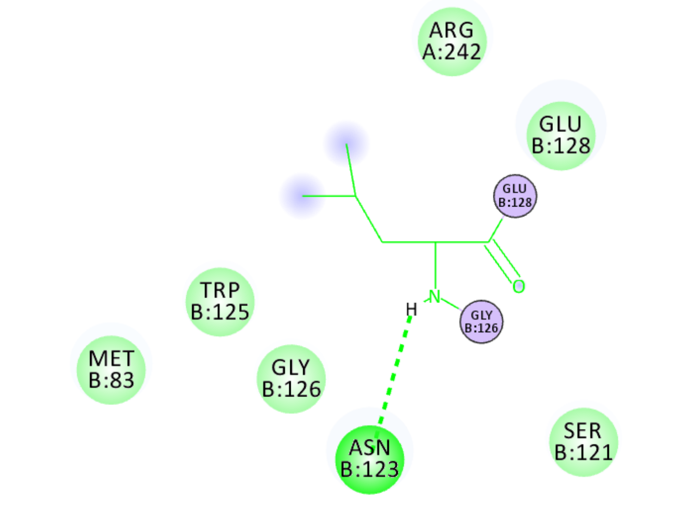


B: Leu127

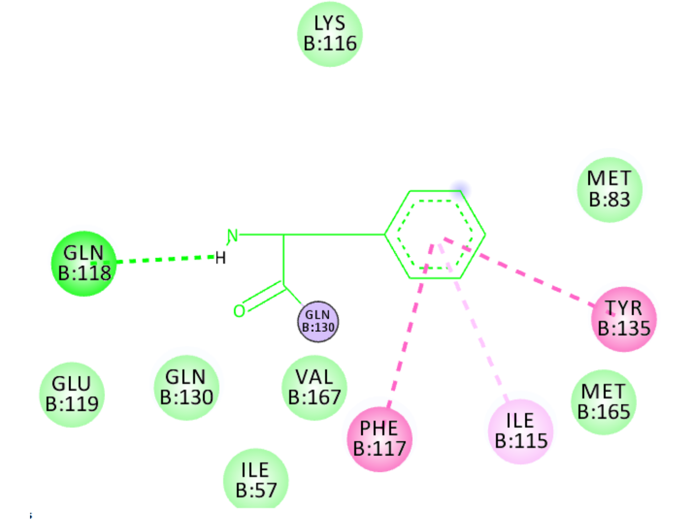


B: Phe129

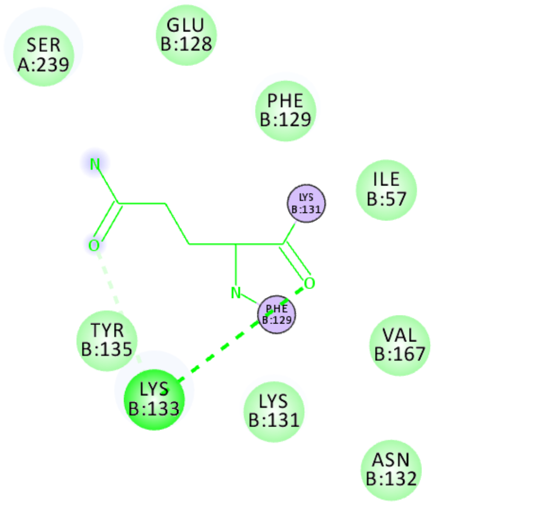


**B: Gln130**

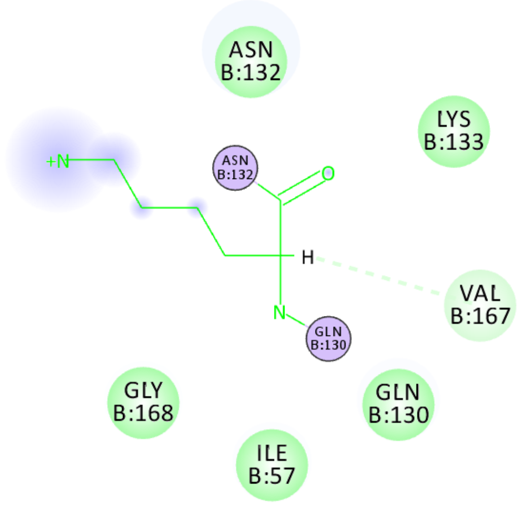


**B: Lys131**

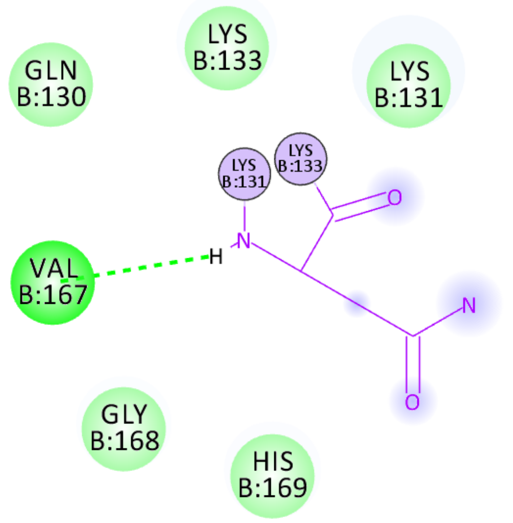


**B: Asn132**

The conventional and carbon hydrogen bonds are shown in bright green/green dashed arrows. The van der Waals are displayed in very faint green lines. Alkyl interactions are colored in light pink dashed line. Covalent interactions are colored in violet. Pi-cation are shown in orange and Pi-Pi T shaped interactions are displayed in bright pink. Amide-Pi stacked is displayed in pink. The interacting amino acids are also colored by the type of interaction. Interacting residue's solvent accessible surface is shown by a blue halo. These color codes of Discovery Studio are maintained here [25].
